# An Objective Detection Method for Tinnitus using Memristors

**DOI:** 10.1101/2025.08.24.671744

**Authors:** Hui Ma, Xiaoyan Ma, Yingying Xuan, Zelin Cao, Lin Li, Kaikai Gao, Mengna Wang, Ruifeng Qiao, Qin Hu, Wentao Yan, Kun Wang, Haoyuan Wang, Longhui Fu, Teng Wu, Chensi Xu, Baiya Li, Juan Hu, Bai Sun, Xiao-jun Li

## Abstract

Approximately 10-20% of the world population experiences tinnitus, which is increased with age and comorbid with frustration, annoyance, anxiety, depression, cognitive dysfunction, insomnia and stress. The underlying mechanisms of tinnitus remain largely unknown and there are no quantifiable biomarkers, making this prevalent disorder a global burden. There are no FDA-approved therapeutics and the current assessment mainly relies on self-reports (e.g., questionnaires), which underscore the need for diagnostic tools for tinnitus. Memristors, nanoscale electronic components mimicking synaptic plasticity, can monitor and control neuronal activity, hold transformative potential for detecting and characterizing tinnitus objectively. Here, we harvested the blood from human and mice with tinnitus, and engineered memristors via magnetron sputtering and made the functional layer with serum. Its core mechanism lies in regulating the reversible formation and rupture of conductive filaments which is formed by utilizing abnormally elevated glutamate concentrations in serum assisted Ag^+^ ions. This serum-based memristor achieves unprecedented discrimination between tinnitus and non-tinnitus individuals. Furthermore, this platform provides mechanistic insights into tinnitus pathogenesis, with potential implications in other neurological disorders.

## 1. Introduction

Tinnitus is an abnormal auditory condition, which characterized by phantom sounds (e.g., buzzing or ringing) in the absence of external stimuli. The prevalence of tinnitus is about 10 to 15% among the adults and as high as 20%-30% in people over 60 years. Tinnitus can also be caused by noise induced hearing loss, presbycusis, Meniere’s disease, virus or bacterial infection, medical conditions such as high blood pressure, otosclerosis, diabetes, and allergies. Approximately 80% of people with tinnitus also reported hearing loss, leading to communication deficits and social isolation.^[1]^ The severity of tinnitus is heterogeneous, ranging from mildly bothersome to extremely disruptive. Chronic tinnitus may exacerbate psychological problems such as anxiety and depression, and correlates with elevated suicide risk.^[2]^ Hair cell or spiral ganglion neuron damage attenuates neural activity transmitted from cochlea to the auditory cortex, this sensory deficit enhanced the gain of the central auditory system in the central auditory system, and this compensatory hyperexcitability is widely hypothesized to be the main pathogenic mechanism of tinnitus.^[3, 4]^

Tinnitus is a subjective symptom, with the pathogenesis and severity vary from person to person. The current diagnosis of tinnitus predominantly depends on the subjective reports of patients, lacking objective criteria to understand and identify the heterogeneity of tinnitus. Despite the clinical utility of cognitive behavioral therapy (CBT) for severe chronic tinnitus,^[1]^ standardized interventions for this auditory neuropathy such as device, pharmaceuticals, or objective diagnostics remain elusive.^[5]^ Pure tone audiometry and tinnitus matching test can quantify the frequency of hearing loss and the loudness of tinnitus, however, it is impossible to elucidate the underlying pathogenesis.^[6]^ Computed tomography (CT) and magnetic resonance imaging (MRI) play effective roles in excluding temporal bone lesions or vestibulocochlear nerve anomalies in tinnitus patients,^[7]^ but they are very limited during the diagnosis of idiopathic tinnitus.^[8]^ Furthermore, there are no tinnitus biomarkers, and the existing tests are deficient to objectively and accurately evaluate this disease. Consequently, advancing objective assessment tools is imperative to refine diagnostic precision, therapeutic targeting, and clinical management.

Memristors, passive circuit elements that functionally emulate neuronal synapses by dynamically modulating resistance via ion migration, retain a relationship between the time integrals of current and voltage across a two-terminal element.^[1, 9–11]^ Memristors achieve resistive switching (RS) behavior similar to neurotransmitter release by electrochemical migration of metal ions to form conductive filaments (CF). Previous studies have showed the signaling efficacy of memristor in bionic sensory systems (e.g., tactile, vision, olfactory systems) (**Table 1**), but there is still a gap in the field of auditory. This study will help to establish a connection between peripheral blood test and central nervous system disorder of tinnitus, creating a new paradigm of neuromorphic electronics in audiological diagnostics, and promoting the development of individualized medical treatment and smart wearable devices.

**Table 1.** Current applications of memristors in medicine.

| Applications | Functions | Ref. |
| --- | --- | --- |
| Tactile | Memristor are utilized to mimicking human thermal nociceptors, with in-situ temperature sensing, internal storage, and computing capabilities. | [12] |
|  | Memristors are utilized to simulate human skin, sense physical signals such as external pressure and temperature. | [13] |
| Vision | Integration of memristors with other devices for modeling biological vision system functions such as depth perception, eyestrain, etc. | [14] |
|  | Memristors are fabricated as opto-electronic memristors (OEMs) and integrated in 1T1OEM arrays for vision processing tasks such as image preprocessing, target tracking, and human motion recognition. | [15] |
| Olfactory | Memristors are fabricated as gas-sensitive resistors for gas sensing, especially for the detection of nitric oxide. | [16] |

In this study, memristors with an Ag/Serum/Ti structure were constructed by employing magnetron sputtering method. Titanium (bottom electrode) ensures interfacial stability and biocompatibility, while silver (top electrode) provides optimal conductivity and electrochemical stability.^[17]^ These devices exhibit excellent resistive switching (RS) window at lower voltage biases (±1.5 V), which are governed by Ohmic conduction and ionic rearrangement as indicated by further fitting the *I-V* curve and modeling of charge transport.^[18]^ Crucially, memristors fabricated with serum from tinnitus patients and noise-induced tinnitus mice display distinct electrical signatures versus controls, enabling specific tinnitus detection. This work pioneers memristor-driven, personalized *in vitro* diagnostics for tinnitus, highlighting their transformative potential in intelligent monitoring in biomedicine.

## 2. Results and Discussion

### 2.1 Establish of noise induced tinnitus mouse model

About 80% of tinnitus patients also reported hearing loss,^[19]^ and noise exposure is one of the popular methods in establishing tinnitus mouse mode. In this study, we firstly constructed noise induced tinnitus mouse model by exposing one of the ear with 16 kHz noise at 116 dB SPL for 45 min and another ear was protected from noise exposure with earplug (**Figure 1**a).^[20]^ Auditory Brainstem Response (ABR), also known as Brainstem Auditory Evoked Potential (BAEP), is a neurophysiological test that evaluates the electrical activity of the auditory pathway from the cochlea to the brainstem in response to sound stimuli. It is widely used to assess hearing function and diagnose neurological disorders in mice.^[21]^ The acoustic trauma by noise exposure was further identified by measuring the threshold of ABR to a click and tone stimuli, including 4 kHz, 8 kHz, 12 kHz, 16 kHz, 24 kHz and 32 kHz. After noise exposure, a significant increase of hearing threshold was observed in 16 kHz and 24 kHz of non-tinnitus mice, and from 8 kHz to 32 kHz in tinnitus mice (Figure 1b-d).

Animals exhibit a natural startle reflex to a sudden loud sound (startle stimulus). A weak, non-startling prepulse (e.g., a tone or gap in noise) presented shortly before the stimulus will reduces the startle response.^[22]^ The gap startle inhibition paradigm is a behavioral experiment that is widely used during the identification of tinnitus in animal models. Its principle relies on the disruption of auditory gap detection caused by phantom auditory perception (tinnitus), which affects the animal’s detection of brief silent intervals in background noise. In the gap startle paradigm, the prepulse is a brief silent gap embedded in continuous background noise. Normal animals can detect this gap, which prevents their startle response to the subsequent loud sound stimulus. Animals with tinnitus possess a constant internal “phantom sound” masking their ability to detect the silent gap in external noise. This was quantified as the gap startle ratio: Gap Startle Ratio=Startle amplitude without gap/Startle amplitude with gap. Higher ratios indicate impaired gap detection, suggesting the presence of tinnitus (Figure 1a). The gap startle ratios did not change significantly across all frequencies in control and non-tinnitus mice before and after noise exposure (Figure 1e-f). A significant increase in the gap startle ratio was observed at 16 kHz in the tinnitus group (Figure 1g).

Repulse Inhibition (PPI) is a neurophysiological phenomenon, in which a weak sensory stimulus (prepulse) presented shortly before a startle-inducing stimulus can reduce the magnitude of the startle reflex. This paradigm is widely used to assess sensorimotor gating and auditory processing in tinnitus. In the pre-pulse inhibition paradigm, a non-startling prepulse (e.g., a tone) delivered 50 ms before the startle stimulus activates inhibitory neural circuits, suppressing the startle reflex. The PPI test was used to exclude severe hearing loss and sensory gating dysfunction in tinnitus mice (Figure 1a).^[23]^ PPI startle ratio was calculated in control, non-tinnitus, and tinnitus groups to exclude potential temporal processing deficits or behavioral problems. No significant differences were observed between control, non-tinnitus, and tinnitus groups across all frequencies before and after noise exposure, suggesting that the tinnitus-related behaviors of mice were not attributed to temporal processing deficits (Figure 1h-j).

**Figure 1.**
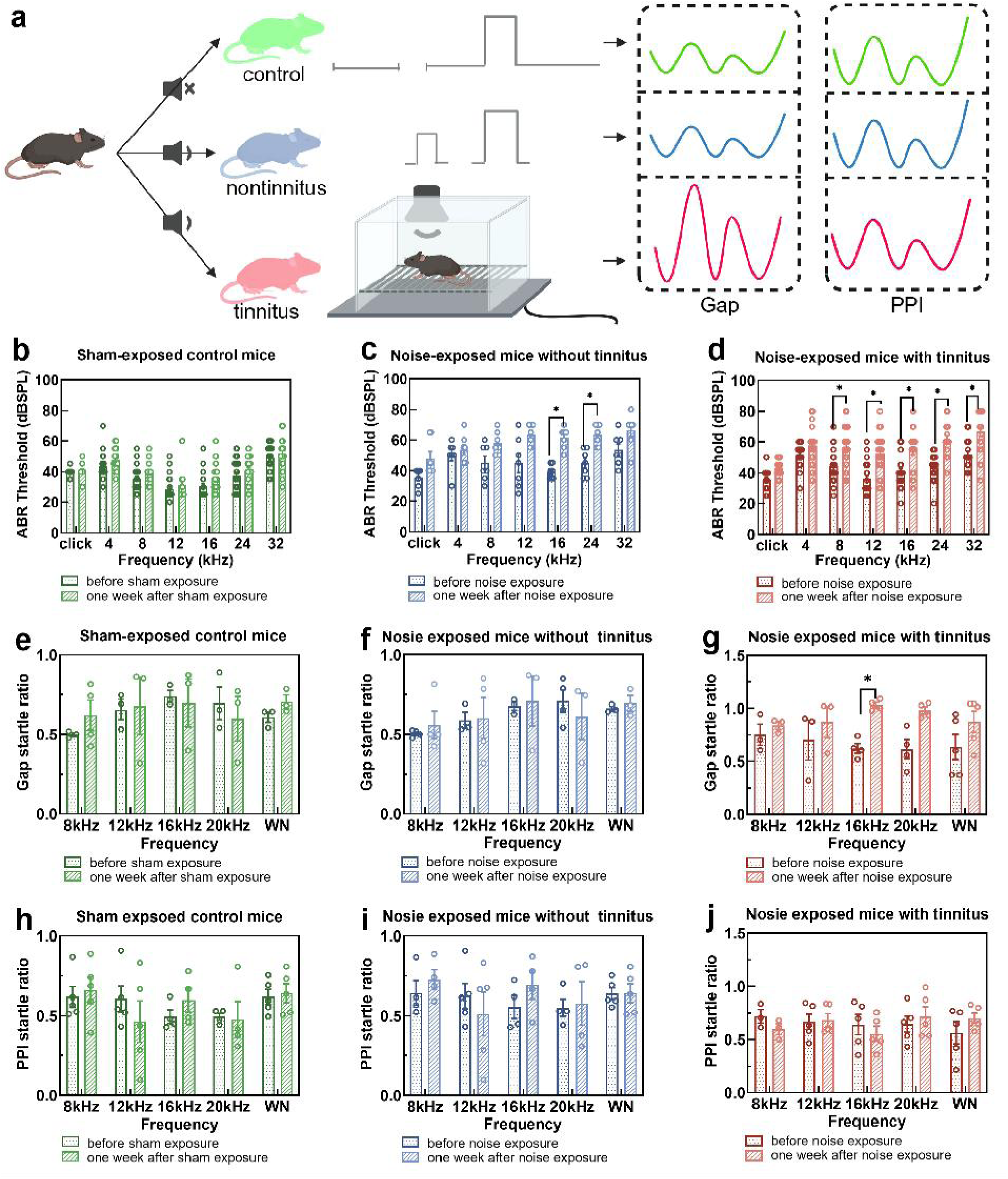
Establish of noise induced tinnitus mouse model. a) Experimental paradigm for gap startle ratio and PPI startle ratio in control, non-tinnitus and tinnitus group (n = 3-5 animals). Pictures were made with BioRender.com released under a Creative Commons Attribution-NonCommercial-NoDerivs 4.0 international license. b) Click- or pure tone-evoked ABR threshold of control mice before and after sham exposure. c) Click- or pure tone-evoked ABR threshold of non-tinnitus mice before and after noise exposure. d) Click- or pure tone-evoked ABR threshold of tinnitus mice before and after noise exposure. e) Gap startle ratio of control mice before and after sham exposure. c) Gap startle ratio of non-tinnitus mice before and after noise exposure. f) Gap startle ratio of tinnitus mice before and after noise exposure. g) Gap startle ratio before and one week after noise exposure. h) The prepulse inhibition (PPI) startle ratio of control mice before and after sham exposure. i) PPI startle ratio of non-tinnitus mice before and after noise exposure. j) PPI startle ratio of tinnitus mice before and after noise exposure. n=3-5 animals, two-tailed, unpaired Student’s t test was used to calculate *p* values. *\*p* < 0.05, where error bars indicate SEM.

### 2.2 Construction of memristor with mouse serum

To investigate the application of memristors in tinnitus diagnosis, blood samples were harvested from non-tinnitus and tinnitus mice, and memristors with an Ag/mouse serum/Ti structure were fabricated (**Figure 2**a). The fabrication protocol entailed spin-coating mouse serum onto Ti substrates, followed by air-drying at ambient temperature. Subsequently, an Ag top electrode was deposited via magnetron sputtering onto the serum film layer as described previously.^[22]^ For electrical characterization, a voltage bias sequence was applied to the Ag top electrode, with the Ti bottom electrode grounded throughout the measurements.

Structural and compositional analyses of the fabricated memristors were conducted using multiple characterization techniques. Scanning electron microscopy (SEM) combined with energy-dispersive X-ray spectroscopy (EDX) was employed to examine the elemental distribution within the serum-based middle layer. Surface SEM-EDX mapping revealed the spatial distribution of oxygen (O), carbon (C), and calcium (Ca) elements in the mouse serum (Figure 2b), with detailed elemental profiles provided in Figure S1 (Supplementary Information). Cross-sectional SEM-EDX analysis further visualized the elemental composition across the device layers, where Ag, C, and Ti were color-coded in red, blue, and green, respectively (Figure 2c). Quantitative elemental distribution data for the cross-section are presented in Figure 2d, in which it clearly reveals that the functional layer is serum.

Spectroscopic and diffraction analyses were performed to characterize the serum material properties. Fourier-transform infrared spectroscopy (FTIR, Figure 2e) and X-ray diffraction (XRD, Figure 2f) provide further insights into the chemical functional groups and crystalline structure of the mouse serum respectively. The high-resolution X-ray photoelectron spectroscopy (XPS) spectrum shows that the most abundant elements in the functional layer are C and O (Figure 2g), and the XPS spectra of O1s and C1s exhibited distinct binding energy peaks corresponding to these elements (Figure 2h-i), confirming their chemical states and bonding configurations within the device. Collectively, these multi-modal characterization of the structural integrity and material composition of Ag/mouse serum/Ti memristor promotes its potential use in tinnitus diagnosis.

**Figure 2.**
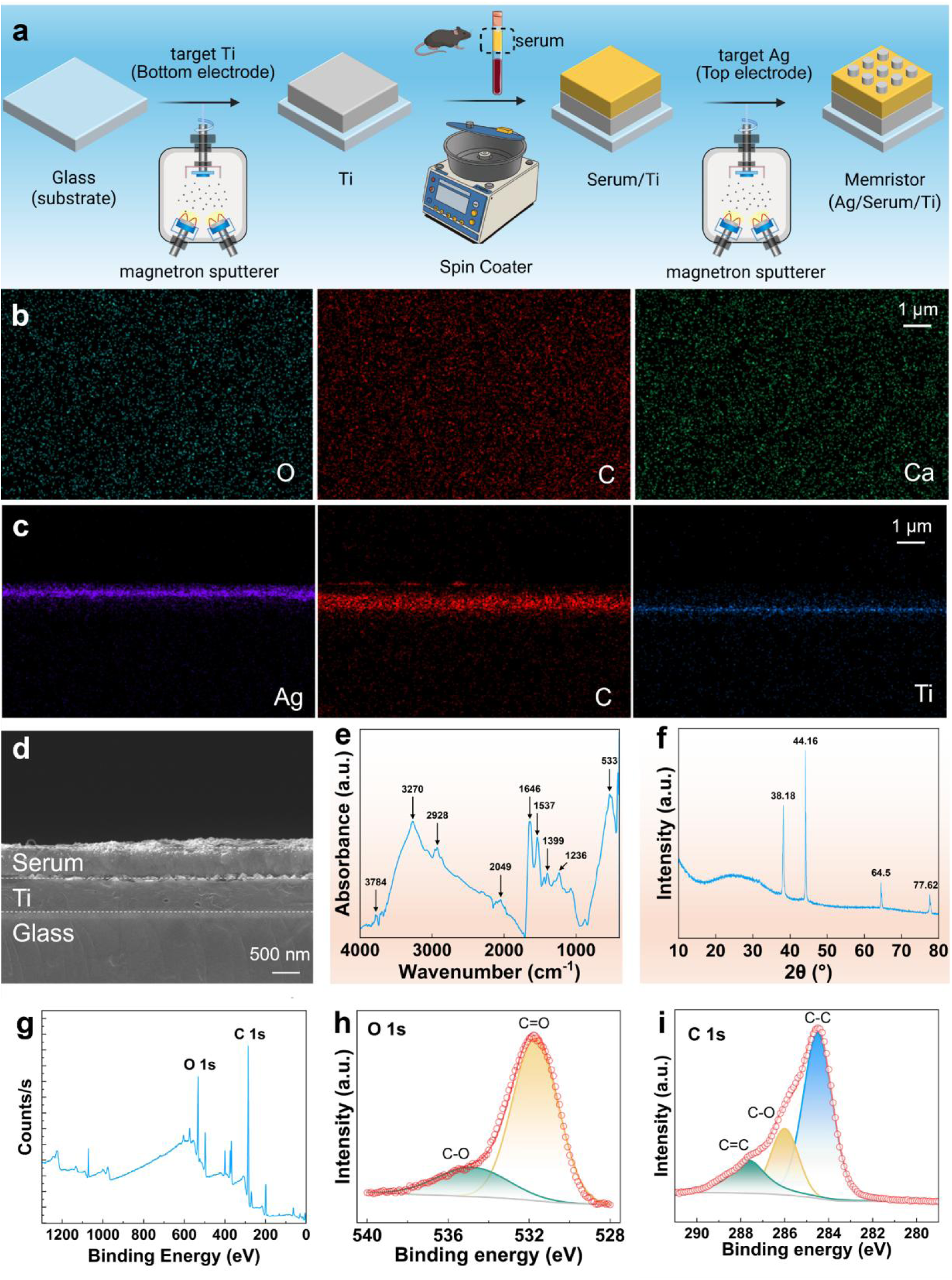
Fabrication and characterization of memristors with an Ag/mouse serum/Ti structure. a) Fabrication process of memristors. Sham or noise exposed mice were scarified and retro-orbital blood were harvested and centrifuged to get fresh serum. Then 100 μL of fresh serum was dropped on an ordinary glass sheet and spin-coating with 500 rpm for 20 s. Finally, Ag was sputtered onto the serum film and used as the top electrode of the memristor. Carton figures were made with BioRender.com released under a Creative Commons Attribution-NonCommercial-NoDerivs 4.0 international license. b) Distribution of different elements on the surface of the memristor under SEM test. c) Distribution of different elements in the cross-section of the memristor under SEM test. d) SEM image of the cross-section of the memristor. e) FT-IR analysis of mouse serum. f) XRD characterization of mouse serum. g) XPS spectrum. h) XPS spectrum of O1s. i) XPS spectrum of C1s.

### 2.3 Diagnosis tinnitus mice with memristor

To establish a discriminative method for tinnitus and non-tinnitus mice, serum samples were harvested from three groups: control, non-tinnitus, and tinnitus mice. Memristive devices were subsequently fabricated using fresh serum, and electrical characterization was conducted by connecting a source meter to the device’s positive and negative electrodes (**Figure 3**a). The current-voltage (*I-V*) characteristic curves were measured within an applied voltage range from -1.5 V to 1.5 V (Figures 3, b-d), where the arrow direction denotes the scanning voltage sequence. It is found that the device initially exist in a high-resistance state (HRS). As the positive voltage was incrementally scanned, the device transitioned from HRS to a low-resistance state (LRS), a process referred to as “SET”. In the negative voltage region, the device reverted from LRS to HRS, which is defined as the “RESET” process. These findings provide conclusive evidence that the memristor device exhibits characteristic bipolar nonvolatile resistive switching behavior, consistent with previous reports.^[23, 24]^ The corresponding logarithmic *I-V* curves are presented in Figures 3e-g. When the voltage window was extended to -2 V to 2 V, the memristive effect of the device was less pronounced compared to the -1.5 V to 1.5 V range (Figure S2).

The device shows time-dependent coordinated curves of voltage and current (Figure 3h), and we performed a comparative analysis of the high and low resistance states across different memristors (Figure 3i), with the inter-group differences in resistance states further emphasized in Figure 3j. When the electrical characteristics of memristors fabricated from different mouse serum were compared under the same voltage bias (-1.5 V to 1.5 V), the *I-V* curves of devices made with control and noise-induced non-tinnitus mouse serum were similar, with no statistical difference between these two groups. In contrast, the *I-V* curves of memristor fabricated from noise-induced tinnitus mouse serum showed a significant deviation from those of non-tinnitus mice. These results strongly imply that the serum compositions of tinnitus and non-tinnitus mice are fundamentally different, and memristor can serve as a viable diagnostic tool for tinnitus using serum.

To comprehensively assess the resistive switching performance of the memristor, additional analyses of its durability and retention characteristics were conducted using Gaussian statistical methods. Figure 3k indicates that the device maintained excellent durability over 50 switching cycles. We further tested 20 more devices to evaluate the variability in HRS and LRS between different devices (Figure S3). The minimal variations in the HRS and LRS values may due to the heterogeneity of the thin-film and the variations of testing parameter setting during fabrication process. The time-dependent retention characteristics of the memristor indicate that the device’s memristive behavior remained stable over time (Figure 3l, m), signifying its reliable nonvolatile storage capabilities. On the basis of these findings, it is further proposed that the developed memristor can be used as an efficient and accurate diagnostic tool for tinnitus in clinical trials.

**Figure 3.**
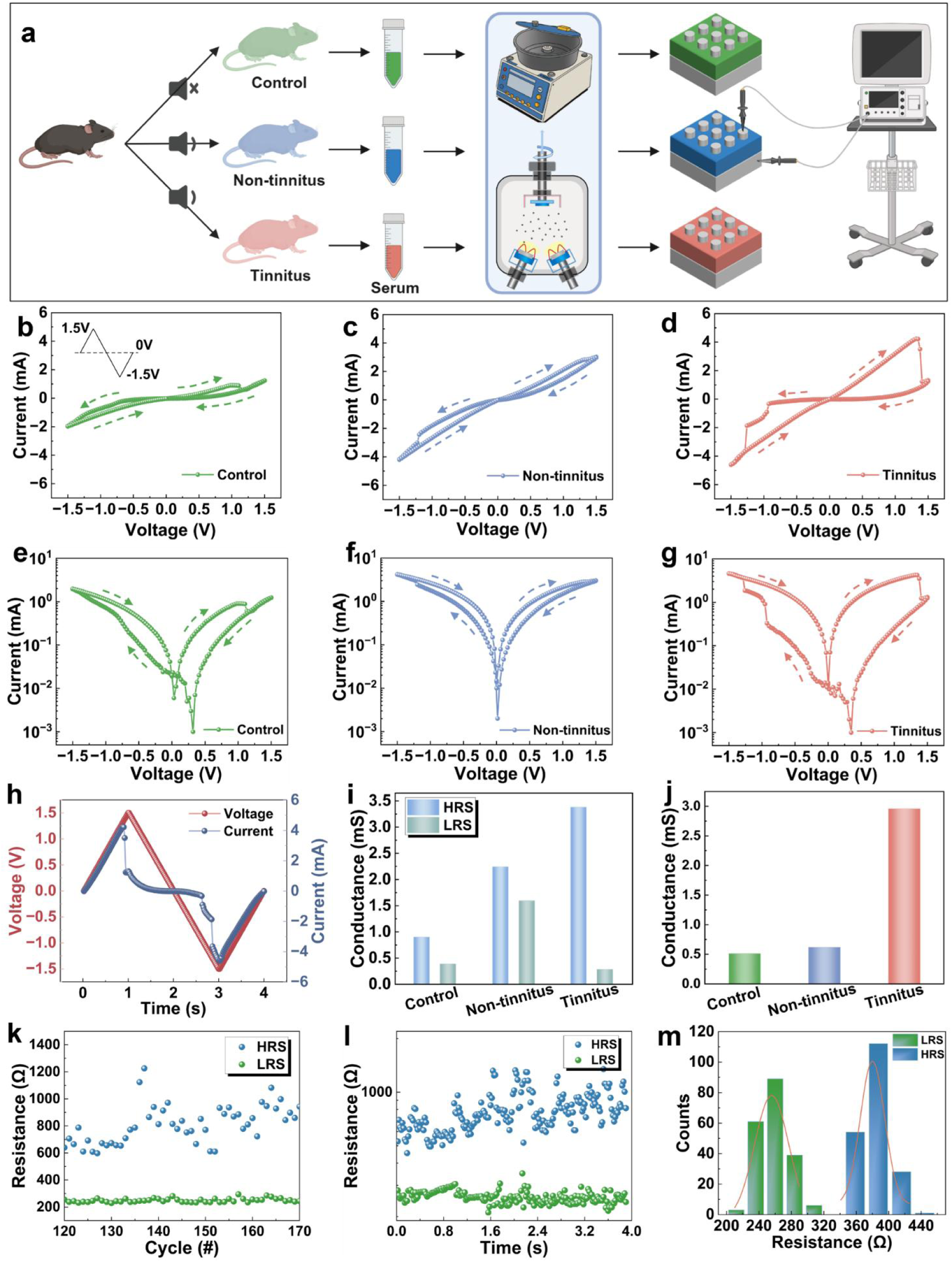
Electrical characteristics of memristor with an Ag/mouse serum/Ti structure. a) Memristors were made with the serum from control, non-tinnitus and tinnitus mice and tested separately. Carton pictures were made with BioRender.com released under a Creative Commons Attribution-NonCommercial-NoDerivs 4.0 international license. b-d) Typical *I-V* curves of memristors. The inset shows the voltage bias parameters applied to the device starting from 0V and cycling between 1.5V and -1.5V. e-g) *I-V* curves in the semi-log scale. h) Time evolution curves of voltage signal and current signal. i) HRS and LRS values of memristors at a voltage of 1.0V. j) The difference between the high and low resistance states in the h plot. k) Endurance test of resistance switching for 50 cycles. l) Retention properties at a read voltage of 1.0 V. m) Gaussian distribution curves of the HRS and LRS, where the red line represents the Gaussian fitting.

### 2.4 The working mechanism model of memristor during tinnitus diagnosis using serum

Memristors transduce signaling similar with synaptic transmition between neurons in the central nervous system (**Figure 4**a).^[24, 25]^ The synapse transmition event invovles release, diffusion, receptor binding of neurotransmitter molecules and unidirectional signal transduction between two neurons. Neurotransmission begins with the arrival of an action potential in the axon terminal and further depolarizing the membrane of the presynaptic terminal, which triggers the release of the neurotransmitters cross the synaptic cleft and bind to receptors on the receiving neuron.^[26, 27]^ The top and bottom electrode in memristor mimics the pre and post synaptic membranes, and the middle layer of memristor is similar with the synaptic cleft. A notable biochemical characteristic in the serum of tinnitus patients is the abnormally elevated level of glutamate.^[28, 29]^ The pH of human serum is between 7.35 and 7.45, and glutamate mainly exists in the form of glutamate ions (Glu⁻). Considering that this device uses a thin film as the functional layer, the interface effect should also be considered as a dominant factor in the working mechanism of the device. When a forward bias voltage (positive voltage) is applied to the memristor, the silver electrode, which acts as the anode, undergoes an oxidation reaction under the electrochemical action and releases silver ions (Ag⁺). At this point, the released Ag⁺ ions undergo a specific coordination reaction with Glu⁻ or glutamate molecules, forming stable silver-glutamate complexes (Figure 4b). These in situ-generated complexes or their subsequent reduction products effectively lower the energy barrier, promoting the reduction deposition and growth of metallic silver at the cathode or adjacent regions, thereby accelerating and stabilizing the formation and bridging of silver-based conductive filaments (CF). The formation of conductive filaments significantly improves the conductivity between the device electrodes, resulting in a significant reduction in the resistance value of the memristor and the device entering a low resistance state (LRS). Similarly, when a negative voltage is applied, the electric field reversal causes the silver-glutamate complex to dissociate, while the anode silver ion supply is interrupted and the filaments are electrochemically dissolved, synergistically blocking the conductive channel and returning the device to a high resistance state (HRS). The high glutamate concentration in tinnitus patients enhances the efficiency of positive pressure filament formation (significantly reducing LRS resistance) and negative pressure filament disruption (significantly increasing HRS resistance), ultimately resulting in a significantly enhanced switching ratio of LRS/HRS, enabling highly sensitive detection of tinnitus through serum. Therefore, elevated glutamate in serum is directly converted into a significant resistance transition from HRS to LRS that can be measured by memristors through the promotion of electrochemically driven silver complexation and filament formation processes, thereby achieving objective detection of tinnitus.

A detailed analysis for the conductivity mechanism of the memristor was conducted via a fitting method to comprehensively understand its excellent applications. Figure 4c shows the *I-V* characteristic curves of the Ag/Serum/Ti-structured memristor under a fixed voltage bias sequence starting from 0V and cycling between 1.5V and -1.5V, with the arrows indicating the order of charge flow. Initially, the device is maintained at LRS, and as the voltage is incremented from 0 to 1.5 V, an abrupt decrease in the current value occurs near 1.2 V, corresponding to the switching of the device to HRS and RESET. Subsequently, the device switches from HRS formation to LRS near -1.2 V, corresponding to the SET process. The alternation of the RESET and SET processes repeatedly gives the device its typical switching characteristics. Further insights into the principles governing LRS and HRS switching are attained through additional fitting of *I-V* characteristic curves and modeling of charge transport. The fitting results indicate that the initial conduction of electrons is mainly Ohmic conduction (Figure 4d). As the electric field strength increases, there is a potential barrier height at the interface between silver and serum. When the applied electric field is large enough, electrons can obtain energy through thermal excitation, cross the potential barrier and enter the conduction band of the insulator. At this point, the main conduction mechanism of electrons shifts to Schottky emission. In this case, the voltage formed by silver ions drives the conductive wire to dominate electron conduction, leading to memory switching from HRS to LRS (Figure 4e, f). When the reverse voltage reaches -1.0V, the conductive channels inside the material become continuous again, and the conduction mechanism shifts back to Ohmic conduction (Figure 4g). With the change of voltage direction, the conduction mechanism shifts to SCLC emission (Figure 4h). Under this mechanism, electrons gradually fill the traps under the action of the electric field, forming a space charge layer, thereby limiting further increase in current. Finally, electrons lack the energy to overcome the potential barrier and become Ohmic conduction during the continuous decrease in reverse voltage (Figure 4i).^[30]^

**Figure 4.**
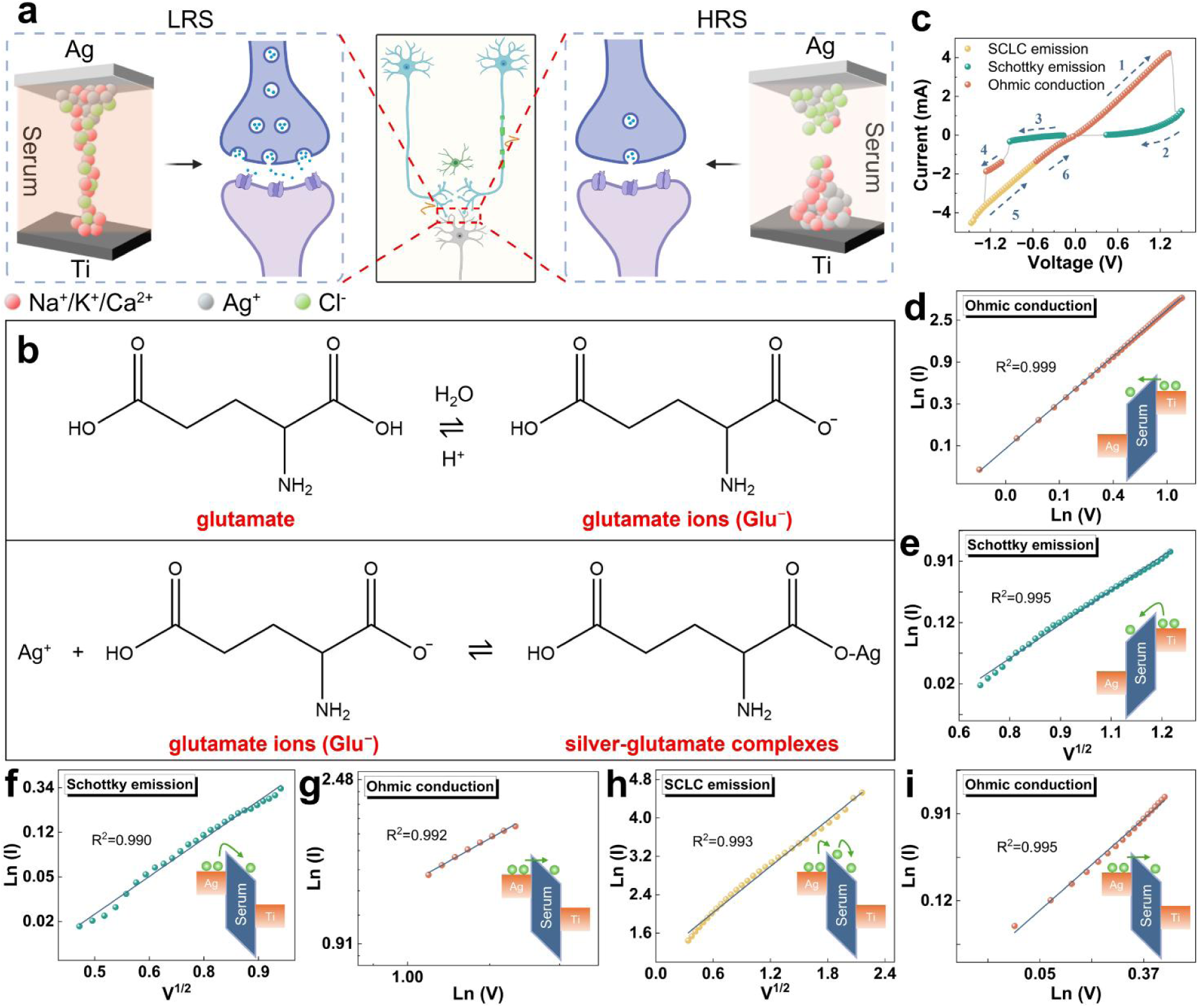
Resistive switching mechanism of the mice blood memristor with an Ag/Serum/Ti structure. a) Sketch of signaling between neurons corresponding to the formation and breakage of conducting filaments during LRS and HRS switching. Carton figures were made with BioRender.com released under a Creative Commons Attribution-NonCommercial-NoDerivs 4.0 international license. b) Signal conduction simulation of memristor with Ag/Serum/Ti structure. The binding interactions between glutamate ions and silver ions. c) Overall diagram of fitted conduction mechanism of the typical *I-V* curve. d-i) Fitting results for the *I-V* curve.

### 2.5 Clinical application of the serum based memristor in tinnitus diagnosis

Based on the above studies, we extend the application of this memristor to the clinic dignosis. Tinnitus is a subjective symptom that is commonly detected by magnetic resonance imaging (MRI), external examination, pure-tone audiometry (PTA), tinnitus handicap inventory (THI), and computed tomography (CT) (**Figure 5**a). Given the subjective heterogeneity of tinnitus and the lack of objective quantitative tools, current diagnosis relies on patient’s medical history and subjective reports such as the Visual Analog Scale (VAS) and the Tinnitus Handicap Inventory (THI), which lacks objective measurement criteria. The Pure Tone Auditory Threshold Test (PTA) and Tinnitus Matching Test can only assess basic audiological parameters such as the frequency of hearing loss and evaluate the loudness of tinnitus, but cannot reflect the intrinsic pathogenesis of tinnitus.^[31]^ In addition, the etiology of tinnitus is complex and includes audiological neurology. Computed tomography (CT) plays an important role in the diagnosis of temporal bone damage in chronic tinnitus, and magnetic resonance imaging (MRI) can be used to evaluate the greater vestibular nerve and rule out the presence of malformations in patients with tinnitus.^[7]^ However, both CT and MRI are limited in the management of idiopathic tinnitus.^[32]^ The lack of tinnitus biomarkers further limits the implementation of precision medicine. Therefore, there is a need to develop more advanced tools and methods to improve the diagnosis, treatment and management of tinnitus. About 80% of tinnitus patients also reported hearing loss, this is consistent with our clinical data, which shows that 7 out of 8 tinnitus patients have hearing loss (**Table 2**) (Figure 5 b-i), with 3 healthy people without tinnitus were used as negative control (Figure 5 j-l). Finally, we performed tinnitus handicap inventory (THI) and found that 62.5% of the tinnitus patients belong to severe disability (Figure 5 m, n).

**Table 2.** The clinical phenotypes of patients with tinnitus.

| No. | Gender | Age | Duration of Symptoms |
| --- | --- | --- | --- |
| 1 | Female | 57 | Left ear: hearing loss for 1 year, accompanied with tinnitus and aural fullness. |
| 2 | Female | 36 | Bilateral tinnitus for 3 months, no hearing loss |
| 3 | Male | 50 | Bilateral tinnitus for 5 years, with hearing loss in the left ear. |
| 4 | Male | 37 | Left ear: tinnitus and hearing loss for 20 years;<br>Right ear: tinnitus for 3 months. |
| 5 | Female | 66 | Left ear: tinnitus and hearing loss for 2 years. |
| 6 | Female | 67 | Hearing loss and tinnitus for 3 years. |
| 7 | Female | 55 | Bilateral hearing loss and tinnitus for 6 years. |
| 8 | Male | 46 | Hearing loss and tinnitus for 6 years. |

**Figure 5.**
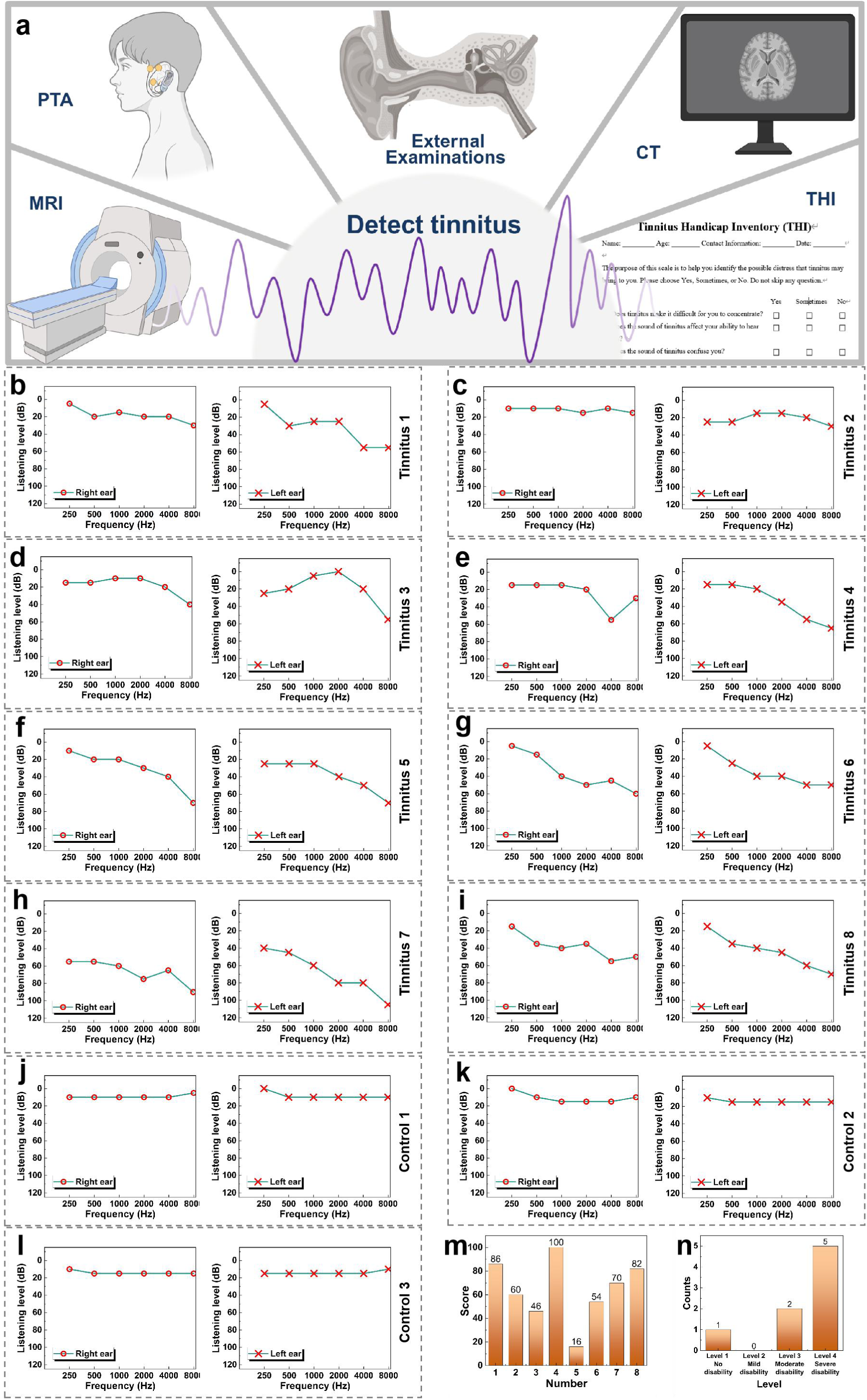
Hearing information of tinnitus patients (n=8). a) Current methods used in detecting tinnitus. Carton figures were made with BioRender.com released under a Creative Commons Attribution-NonCommercial-NoDerivs 4.0 international license. b-i) Pure-tone audiometry (PTA) audiograms of 8 tinnitus patients in the left and right ears. j-l) Pure-tone audiometry (PTA) audiograms of 3 healthy people without tinnitus in the left and right ears. m) Results of test scores of Tinnitus Handicap Inventory (THI) of 8 tinnitus patients. n) Statistics of test grades of Tinnitus Handicap Inventory (THI) of 8 tinnitus patients.

Next, we fabricated the memristors based on serum of tinnitus patients (**Figure 6**a). We analyzed the durability and retention characteristics of the memristor and found that the device maintained excellent durability over 60 switching cycles (Figure 6b). The variations in the HRS and LRS values may due to the heterogeneity of the thin-film and the variations of parameter setting during fabrication process. Then we tested the *I-V* curve under different scan rates, and found that the *I-V* curve of the memristor was significantly different under different scan rates (Figure 6c), we set the scan rate as 0.02s in the following tests. It can be observed that the time-dependent coordinated curves of voltage and current within the device (Figure 6d). Likewise, we tested *I-V* curves of memristors fabricated from serum of normal human (control group) and tinnitus patients (experimental group) over different voltage ranges (Figure 6e, f). On the basis of these findings, it is further proposed that the developed memristor can be used as an efficient and accurate diagnostic tool for tinnitus in clinical trials. The switching ratio (R_off_/R_on_) of memristors in the control group and tinnitus group was compared within the same voltage range (-1V to 1V) (Figure 6g). The experimental results showed that the switching ratio of the control group was only about 1.25, while the tinnitus group was as high as about 7.5. The switching ratio of the control group was significantly smaller than that of the tinnitus group, which was consistent with the results in the mouse model and further supported the working mechanism described above.

**Figure 6.**
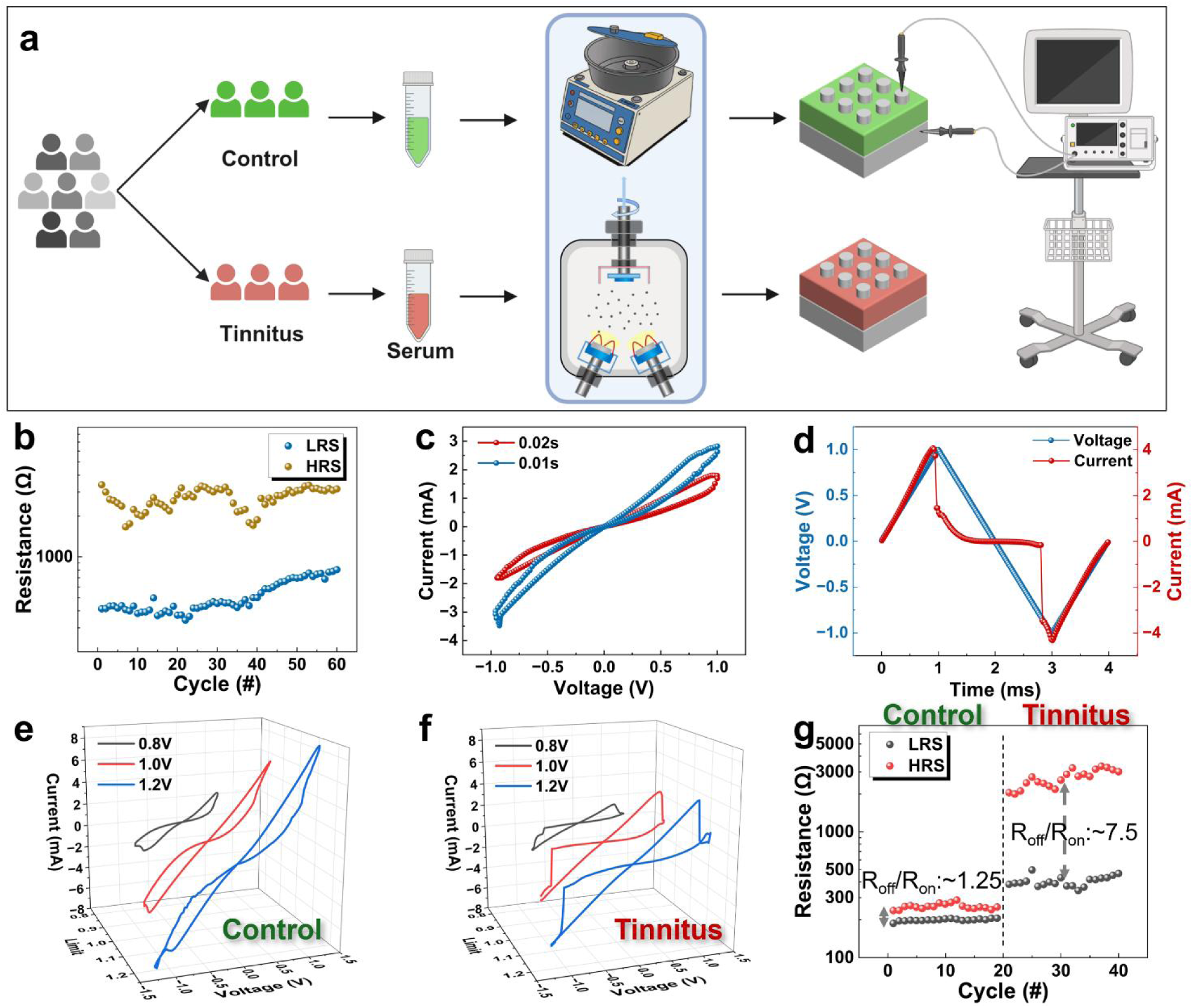
Electrical characteristics of memristor and comparison of electrical properties of memristors fabricated with the blood of healthy and tinnitus patients. a) Experimental scheme of memristors constructed with different serum from human. Carton figures were made with BioRender.com released under a Creative Commons Attribution-NonCommercial-NoDerivs 4.0 international license. b) The endurance test of 60 cycles. c) Characterization of electrical performance curves at different scan rates. d) Time evolution curves of voltage and current signal. e) *I-V* curves of memristors fabricated from serum of healthy people (control group) over different voltage ranges. f) *I-V* curves of memristors fabricated from serum of tinnitus patients (experimental group) over different voltage ranges. g) Comparison of the switching ratio (R_off_/R_on_) of memristors in the control group and tinnitus group.

### 2.6 The future of memristor in tinnitus diagnosis

Tinnitus is recognized globally as a symptom of an underlying health condition. It often accompanies with other disease, including hearing loss and other neurological disorders. Early and accurate diagnosis is not only crucial for timely therapeutic interventions but also for preventing the progression of associated complications. In the context of this study, we explored the potential of memristors as a novel diagnostic modality for tinnitus. Through a series of experiments with serum from human and mice, we were able to fabricate memristors with an Ag/Serum/Ti architecture. Our electrical characterization results, which included detailed analyses of current-voltage (*I-V*) characteristics, logarithmic *I-V* curves, and time - dependent voltage - current relationships, clearly demonstrated that these memristors could accurately distinguish tinnitus human or mice from non-tinnitus control. The characteristic bipolar nonvolatile resistive switching behavior of the memristors, coupled with the significant differences in *I-V* curves between the two groups, provide strong evidence for the diagnostic utility of this approach.

Nevertheless, the translation of this memristor-based diagnostic method from bench to bedside is currently hampered by some intricate nature. To address these challenges and realize the full potential of our memristor-based diagnostic concept, we envision the development of a next-generation diagnostic test strip. This innovative device will be designed based on the fundamental principles of memristor operation, integrating key elements of resistive switching behavior. By leveraging advancements in microfabrication, nanomaterials, and microfluidics, we aim to create a portable, cost-effective, and user-friendly diagnostic tool, as illustrated in **Figure 7**. The test strip will require minimal sample, enable rapid on-site testing, and have the potential to be widely used in various clinical trials from primary care facilities to specialized audiology clinics. This technological breakthrough marks a significant advancement in the diagnosis of tinnitus, which will greatly improve access to timely and accurate medical services for patients suffering from tinnitus.

**Figure 7.**
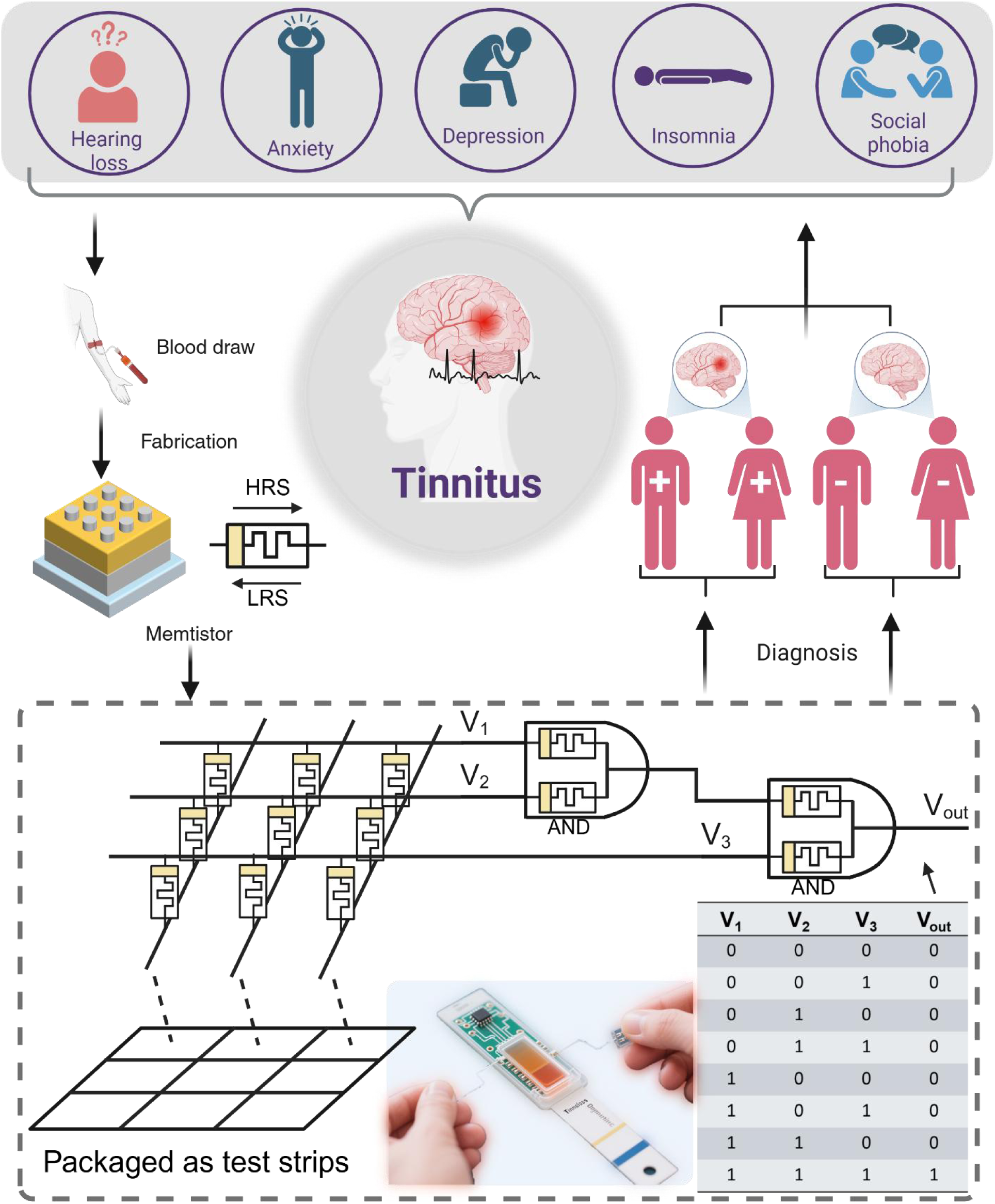
Process of tinnitus detection using a memristor: blood is drawn from tinnitus patients and tested with strips, which are based on a memory array. Patients will be diagnosed as tinnitus only when three voltages are all positive. Then further treatment can be given differentially based on our diagnosis results. Figure was made with BioRender.com released under a Creative Commons Attribution-NonCommercial-NoDerivs 4.0 international license.

## 3. Conclusion

In this study, a biological memristor with Ag/Serum/Ti structure was fabricated by using the resum from noise induced tinnitus mice and tinnitus patientsrespectively. Our results shows that the device exhibited significant RS characteristics at lower voltage biases sequence, and significant differences in electrical characteristics such as *I-V* curves and high/low resistance states among tinnitus and non-tinnitus groups. These findings suggest that memristors can effectively differentiate between tinnitus and non-tinnitus individuals, thus providing a novel and objective method for tinnitus detection. Furthermore, we designed a memristor logic circuit that is expected to be encapsulated into a test strip, which holds great potential in the objective diagnosis of tinnitus. Circuit simulation results validated the effectiveness of the device in distinguishing the presence or absence of tinnitus both in human and mice. These results make the memristor compatible with logic circuits and medical monitoring, promoting early detection of tinnitus and memristor-based biomedical applications.

## 4. Experimental Section

### Mice

C57BL/6 mice at postnatal day 30 (P30) were used for establishing tinnitus mouse model. Experimental protocols were approved by Xi’an Jiaotong University Animal Care and Use Committees protocol (No. XJTUAE2025-198). Both males and females were used in our experiments.

### Auditory Brainstem Response (ABR)

ABR was assessed to evaluate hearing threshold shifts in the animals.^[33]^ Briefly, the mice were anesthetized with 1% to 1.5% isoflurane and placed in a soundproof chamber. Acoustic stimuli with a 0.5-ms rise/fall were directly delivered to the external auditory canal using a TDT RZ6 instrument (Tucker Davis Technologies). Needle electrodes (Rochester Elektro Medical, Lutz, FL, USA), comprising reference, grounding, and active electrodes, were subcutaneously positioned over the head before testing. After averaging 512 times and filtering from 100 to 3 kHz, the amplified responses from the active electrodes were recorded. The intensity of the stimuli was decreased from 90 dB in 10-dB SPL interval for both click and different tone frequencies (4, 8, 12,16, 24, and 32 kHz). The ABR threshold criterion was the lowest intensity at which the repeatable.

### Establish of Tinnitus Mouse Model with Noise Exposure

The animals were anesthetized with 1% to 1.5% isoflurane and fitted with foam earplugs (OHRFRIEDEN, Wehrheim, Germany) in the right ear before being placed in a soundproof chamber equipped with a sound delivery system. A speaker connected to an amplifier was positioned 10cm before the head. 16 kHz continuous noise was delivered for 45min at 116 dB SPL. We identified mice with or without tinnitus after noise exposure by detecting gap and PPI startle ratio.^[20]^

### Gap Detection

We used white noise and narrow bandpass sound with a 1-kHz bandwidth centered at 8, 12, 16 and 20 kHz presented at 70 dB SPL. A 50-ms sound gap was introduced 130 ms before the startle stimulus. Startle response represents the time course of the downward force (presented as arbitrary units, AU) that the mice apply onto the platform in response to the startle stimulus. Gap detection was evaluated by the gap startle ratio, which is the ratio of the peak-to-peak value of the startle waveform in trials with gap over the peak-to-peak amplitude of the startle waveform in trials in the absence of the gap.^[34]^

### Prepulse Inhibition (PPI)

Startle ratio was calculated for the control, non-tinnitus, and tinnitus groups. PPI was tested in a quiet background, and a 50-ms non-startling sound (prepulse) of similar intensity to the background sound used in the gap detection test was presented 130 ms before the startle stimulus. Prepulse inhibition was evaluated by PPI startle ratio, which is the ratio of the peak-to-peak value of the startle waveform in trials with prepulse over the peak-to-peak value of the startle waveform in startle only trial.^[34]^

### Blood Collection

For mouse serum, the animals were anesthetized with 1% to 1.5% isoflurane, and then the blood was collected from mouse retro-orbital using anticoagulant-treated capillary tubes (EDTA/heparinized), followed by centrifugation at 1000 rpm for 10 minutes. The upper serum layer was aspirated using a sterile micropipette, avoiding contamination from the intermediate leukocyte buffy coat or erythrocyte pellet. Similarly, about 5ml of human blood was harvested to collect serum. Both mouse and human serum were used to make Memristors as described previously.^[35]^

### Device Fabrication

In this study, the memristor with Ag/Serum/Ti structure was fabricated using a typical double-ended structure. Specifically, we used magnetron sputtering equipment to sputter titanium (Ti) metal onto an ordinary glass sheet (2×2 cm) as the bottom electrode of the device. The sputtering process parameters for Ti mainly include a chamber pressure of 1.2 pa, a DC voltage of 70 w for 20 min, and a sample tray rotation speed of 3 rpm. Then, we dropped 100 μL of fresh serum onto the glass sheet sputtered with titanium metal (Ti). Then the serum was applied to the substrate by spin-coating (500 rpm, 20 s). After the serum was naturally coagulated, metallic silver (Ag) sputtered on the serum film using a magnetron sputtering apparatus was used as the top electrode of the device. The essential sputtering parameters included a chamber pressure of 1.2 pa, a DC voltage of 70 W, 15 min and a sample tray speed of 3 rpm. Here, a typical Ag/Serum/Ti/Glass-structure memristor was fabricated as previously described.^[36]^

### Device Characterization

The electrical characteristics of the devices were tested using an electrochemical workstation equipped with a source meter (B2901B, Keysight, Beijing, China). The cross-sectional morphology and elemental distribution on the surface of the memristor with Ag/Serum/Ti structure were observed by scanning electron microscopy. The chemical composition of the serum in the dielectric layer of the device was analyzed by FT-IR, XRD characterization and XPS spectrum. Finally, the construction and simulation of the logic circuit based on the blood memristor was realized by Simulink unit in MATLAB.

## Supporting Information

Supporting Information is available from the Wiley Online Library or from the author.

## Acknowledgements

H. Ma, X. Ma and Y. Xuan contributed equally to this work. This work has been supported by National Natural Science Foundation of China (52375575), Natural Science Basic Research Plan in Shaanxi Province of China (Program No.2024JC-ZDXM-43), National Natural Science Fund for Excellent Young Scientists Fund Program (Program No. GYKP045), and the Fundamental Research funds for the Central Universities (to X.-J. Li), the basic research project of Xi’an Jiaotong University (xzd012024058; xtr072024022; xtr062025002; xtr052025010). The authors extend the gratitude from Shiyanjia Lab (www.shiyanjia.com) for providing invaluable assistance with the SEM, XPS and XRD analysis. Parts of the Figures were drawn by using pictures from Servier Medical Art. Servier Medical Art by Servier is licensed under a Creative Commons Attribution 3.0 Unported License (https://creativecommons.org/licenses/by/3.0/).

## Data Availability Statement

The datasets that support the findings of this study are available from the corresponding authors upon reasonable request.

---

In this study, Ag/Serum/Ti memristors were fabricated using serum from control, tinnitus, and non-tinnitus mice groups, which shows significant resistive switching (RS) characteristics and resistance states under low voltage region. This result indicates that the memristors can effectively distinguish tinnitus conditions, offering a novel detection method. Further, a memristor logic circuit was designed for potential use in tinnitus test strips. Finally, the simulation results validate its effectiveness in detecting tinnitus, highlighting its promise for biomedical applications.

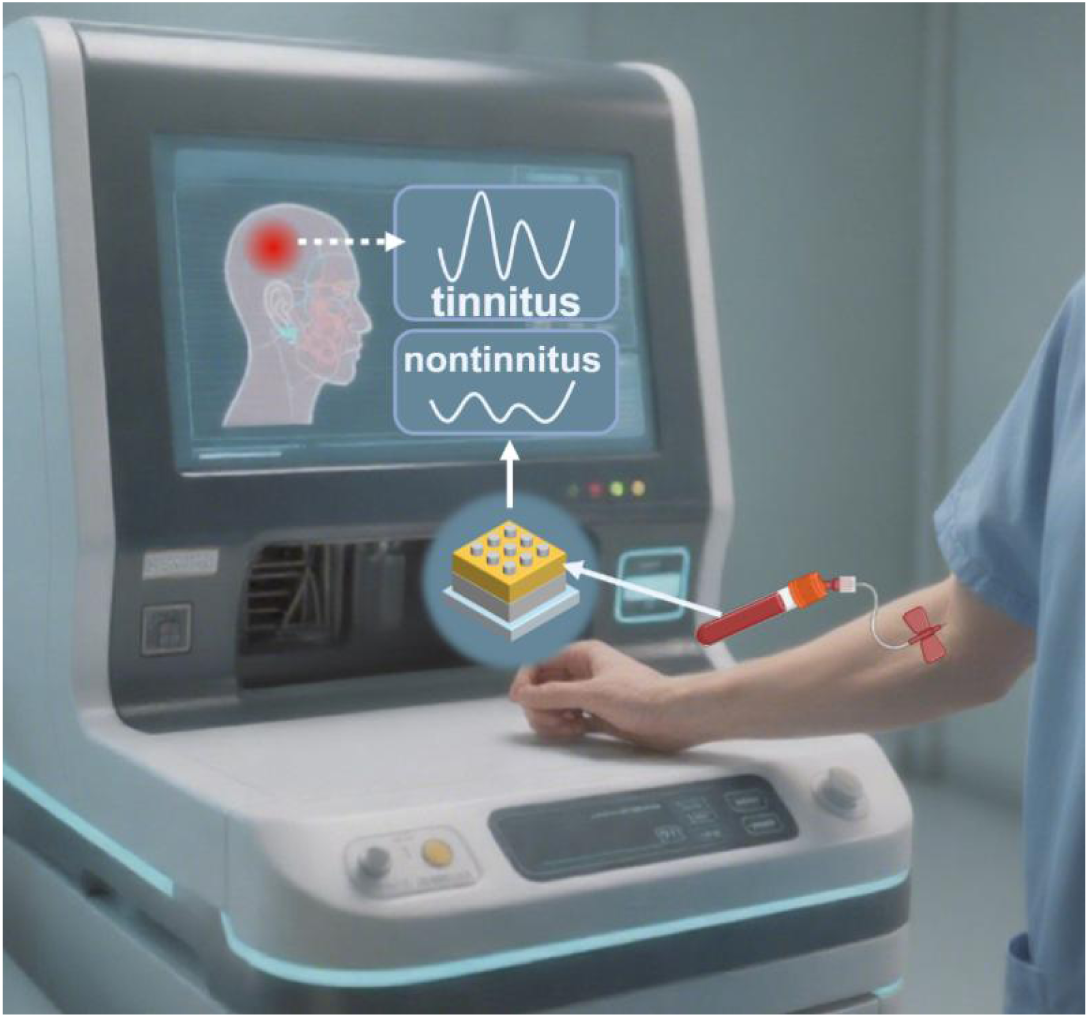

