## Supplementary material for "An Objective Detection Method for Tinnitus using Memristors": 6

**ADVANCED MATERIALS**

### Supporting Information

#### An Objective Detection Method for Tinnitus using Memristors

*Hui Ma, Xiaoyan Ma, Yingying Xuan, Zelin Cao, Lin Li, Kaikai Gao, Mengna Wang, Ruifeng Qiao, Qin Hu, Wentao Yan, Kun Wang, Haoyuan Wang, Longhui Fu, Teng Wu, Chensi Xu, Baiya Li, Juan Hu*, Bai Sun*, and Xiao-jun Li**

H. Ma, Y. Xuan, Z. Cao, L. Li, K. Gao, M. Wang, R. Qiao, Q. Hu, W. Yan, K. Wang, H. Wang, Prof. B. Sun, Prof. X. Li

Frontier Institute of Science and Technology, and Interdisciplinary Research Center of Frontier Science and Technology, Xi'an Jiaotong University, Xi'an, Shaanxi 710049, China

 (X.L.); (B.S.).

X. Ma, B. Li, Prof. X. Li

Department of Otorhinolaryngology Head and Neck Surgery, The First Affiliated Hospital of Xi'an Jiaotong University, Xi'an, Shaanxi 710061, China

C. Xu, Prof. J. Hu

Department of Otorhinolaryngology Head and Neck Surgery, The Second Affiliated Hospital of Xi'an Jiaotong University, Xi'an, Shaanxi 710061, China

 (J.H.).

L. Fu, T. Wu

Department of Neurosurgery, The Second Affiliated Hospital of Xi'an Jiaotong University, Xi'an, Shaanxi 710004, China

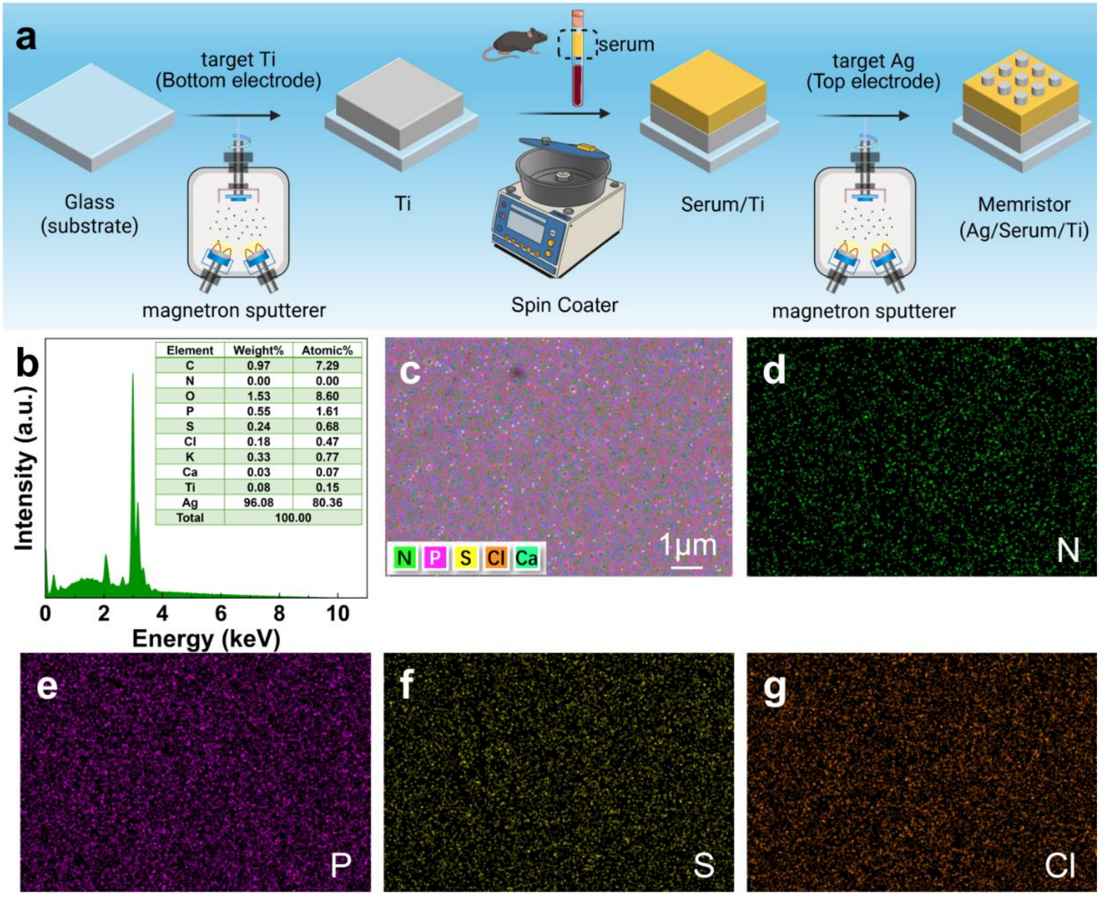


**Figure S1**. Fabrication and microscopic characterization of memristors with Ag/mouse serum/Ti structures. a) Fabrication process of memristors. Created with BioRender.com released under a Creative Commons Attribution-NonCommercial-NoDerivs 4.0 international license. b) Energy dispersive spectrometer (EDS) peak spectrum. c) The energy dispersive spectrometer (EDS) spectrum. d-g) EDS element mapping images of N, P, S and Cl, respectively.

**Supplement Text Note 1:**

Sham or noise exposed mice were scarified and retro-orbital blood was harvested and centrifuged to get fresh serum. Then 100 μL of fresh serum was dropped on a glass sheet and spin-coating with 500 rpm for 20 s. Finally, Ag was sputtered on the serum film and used as the top electrode of the memristor. An Ag/mouse serum/Ti memristor was established. The thickness distribution is uniform, and the boundary is clearly visible. The energy dispersive spectrometer (EDS) peak spectrum is shown in Figure S1a, where the elements O, C, and P can be clearly visualized with atom percentages of 8.6 %, 7.29 %, and 1.61 %, respectively. The serum functional layers were analyzed using SEM-EDX elemental mapping, as shown in Figure S1b. It can be observed that N, P, S, Cl and Ca elements are uniformly distributed (Figure S1d-g), consistent with the arrangement in the EDX spectrum.

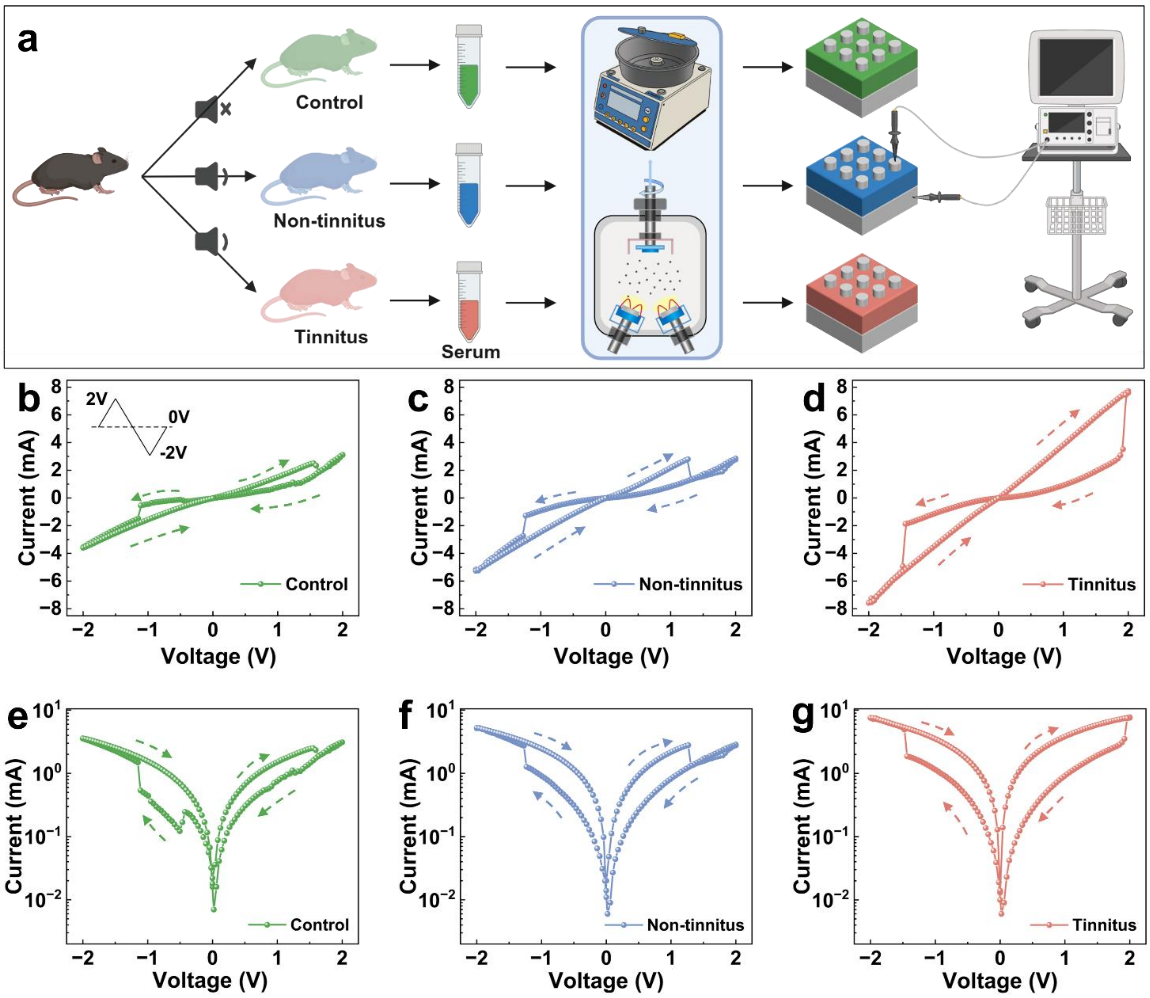


**Figure S2**. Electrical characteristics of memristor with an Ag/mouse serum/Ti structure. a) Testing procedure of memristors prepared using three groups of mice serums separately. Created with BioRender.com released under a Creative Commons Attribution-NonCommercial-NoDerivs 4.0 international license. b-d) Typical *I-V* curves of memristors using serum from control, non-tinnitus, and tinnitus mice. The inset shows the voltage bias parameters applied to the device starting from 0V and cycling between 2V and -2V. e-g) I–V curves in the semi-log scale.

**Supplement Text Note 2:**

In order to determine the optimal scanning voltage, the typical *I-V* characteristic curves of the memristor between 2V and -2V were characterized, as shown in **Figure S2**b-g. The device does not exhibit as much of a memristor effect in the voltage window from -2 V to 2 V as it does from -1.5 V to 1.5 V. However, the tinnitus group has a larger switching window than the non-tinnitus group and the control group.

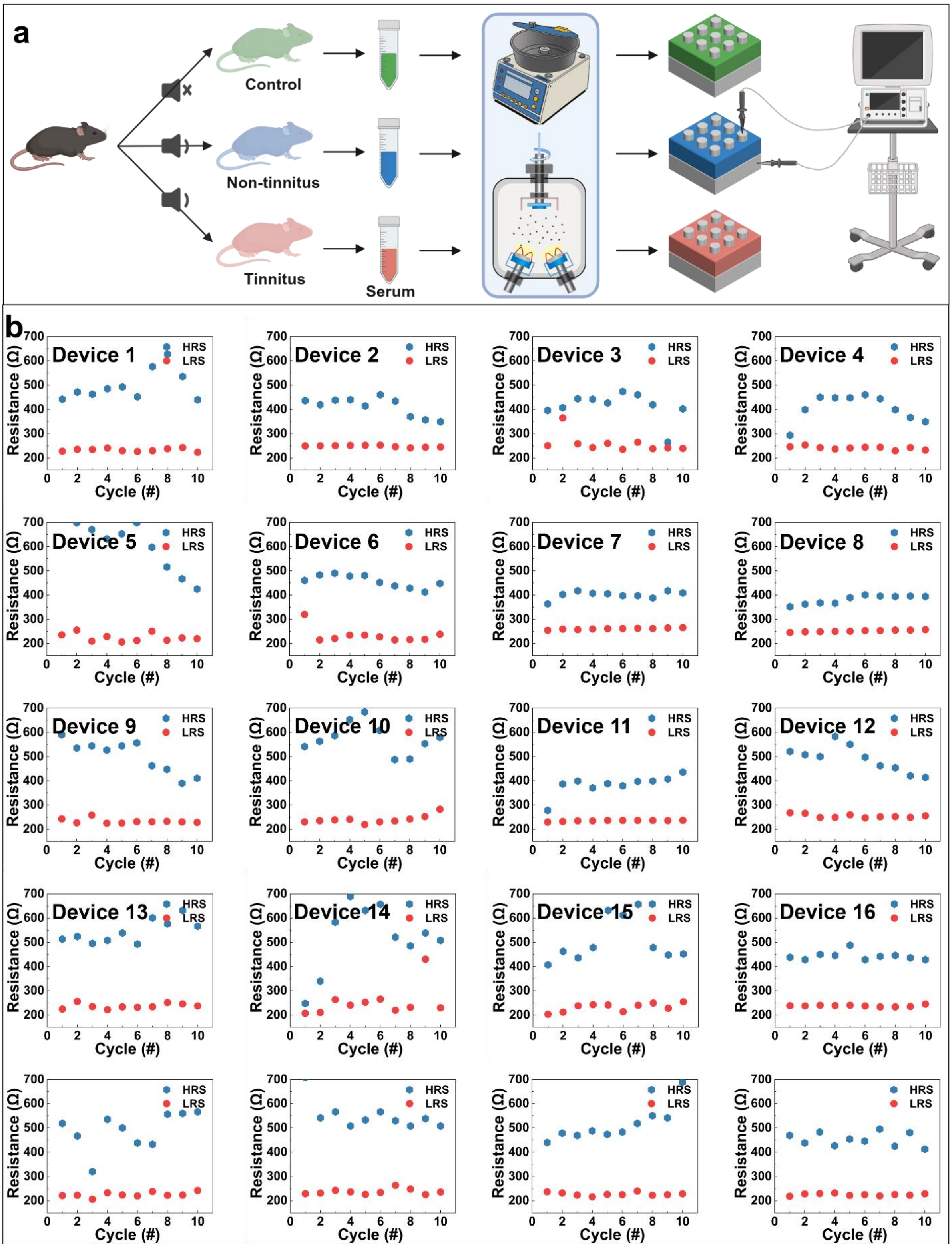


**Figure S3**. a) Testing procedure of memristors prepared using serum from control, non-tinnitus and tinnitus mice separately. Created with BioRender.com released under a Creative Commons Attribution-NonCommercial-NoDerivs 4.0 international license. b) The HRS and LRS of 20 memristors using serum from tinnitus mice under the same voltage sweep range.

**Supplement Text Note 3:**

As shown in **Figure S3**b, we tested 20 devices to investigate the differences between the high and low resistance states of different devices using serum from tinnitus mice. We found that the HRS and LRS of the 20 devices are close with each other. Some of the variations come from the uniformity of the serum films and the fabrication process between devices.

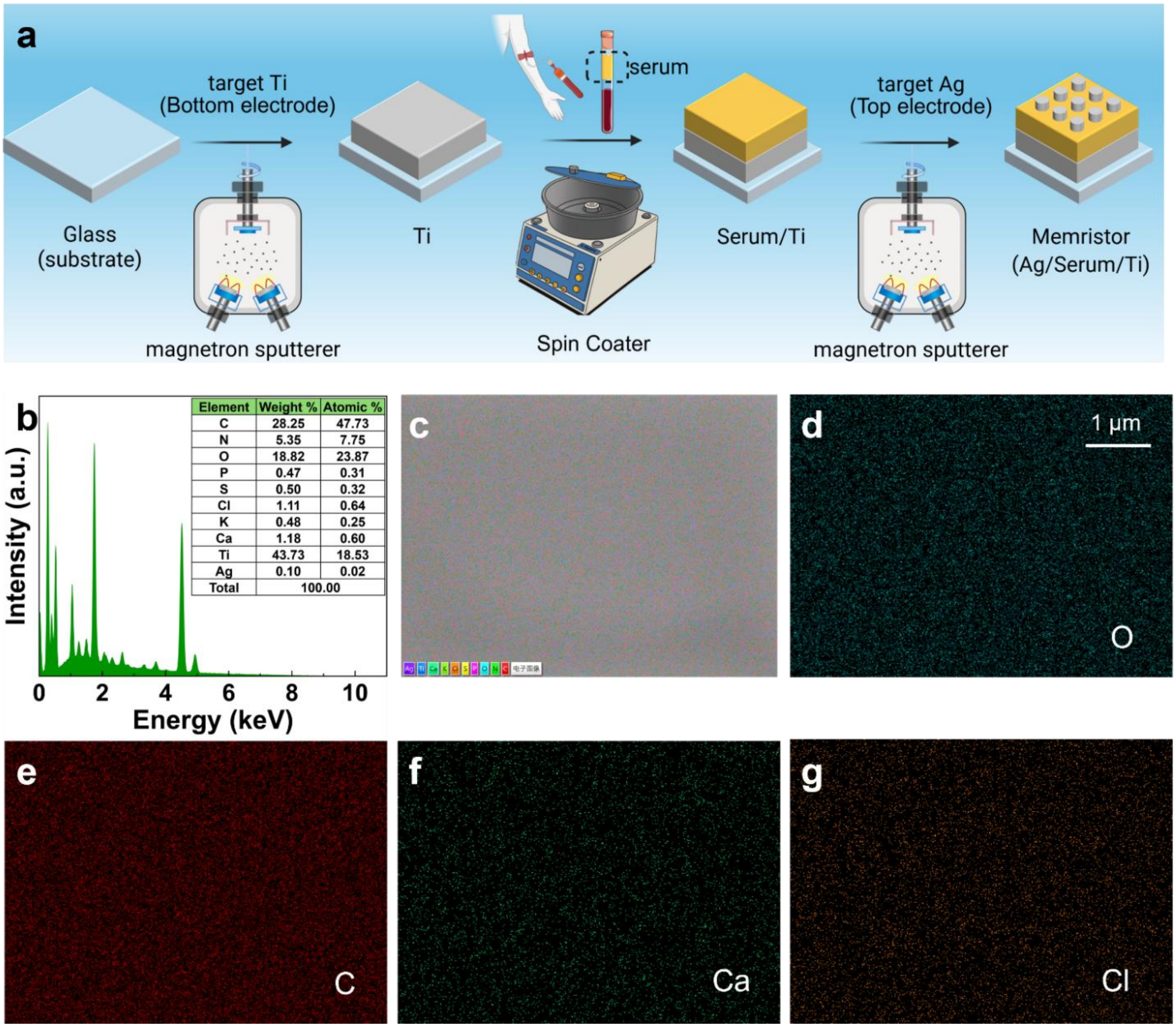


**Figure S4**. Establish and microscopic characteristics analysis of memristor with Ag/human serum/Ti structure. a) Fabrication process of memristor with Ag/human serum/Ti structures. Created with BioRender.com released under a Creative Commons Attribution-NonCommercial-NoDerivs 4.0 international license. b) Energy dispersive spectrometer (EDS) peak spectrum. c) The energy dispersive spectrometer (EDS) spectrum. d-g) EDS element mapping images of O, C, Ca and Cl, respectively.

**Supplement Text Note 4:**

Blood from healthy and tinnitus patients were harvested and centrifuged to get fresh serum. Then 100 μL of fresh serum was dropped on a glass sheet and spin-coating with 500 rpm for 20 s. Finally, Ag was sputtered on the serum film and used as the top electrode of the memristor (**Figure S4**a). The energy dispersive spectrometer (EDS) peak spectrum is shown in Figure S4b, where the elements O, C, and P can be clearly visualized with atom percentages of 23.87 %, 47.73 %, and 0.31 %, respectively. An Ag/human serum /Ti memristor was constructed. The thickness distribution is uniform, and the boundary is clearly visible. The serum functional layers are analyzed using SEM-EDX elemental mapping, as shown in Figure S4c. It can be observed that O, C, Ca and Cl elements are uniformly distributed, consistent with the arrangement in the EDX spectrum (Figure S4 d-g).

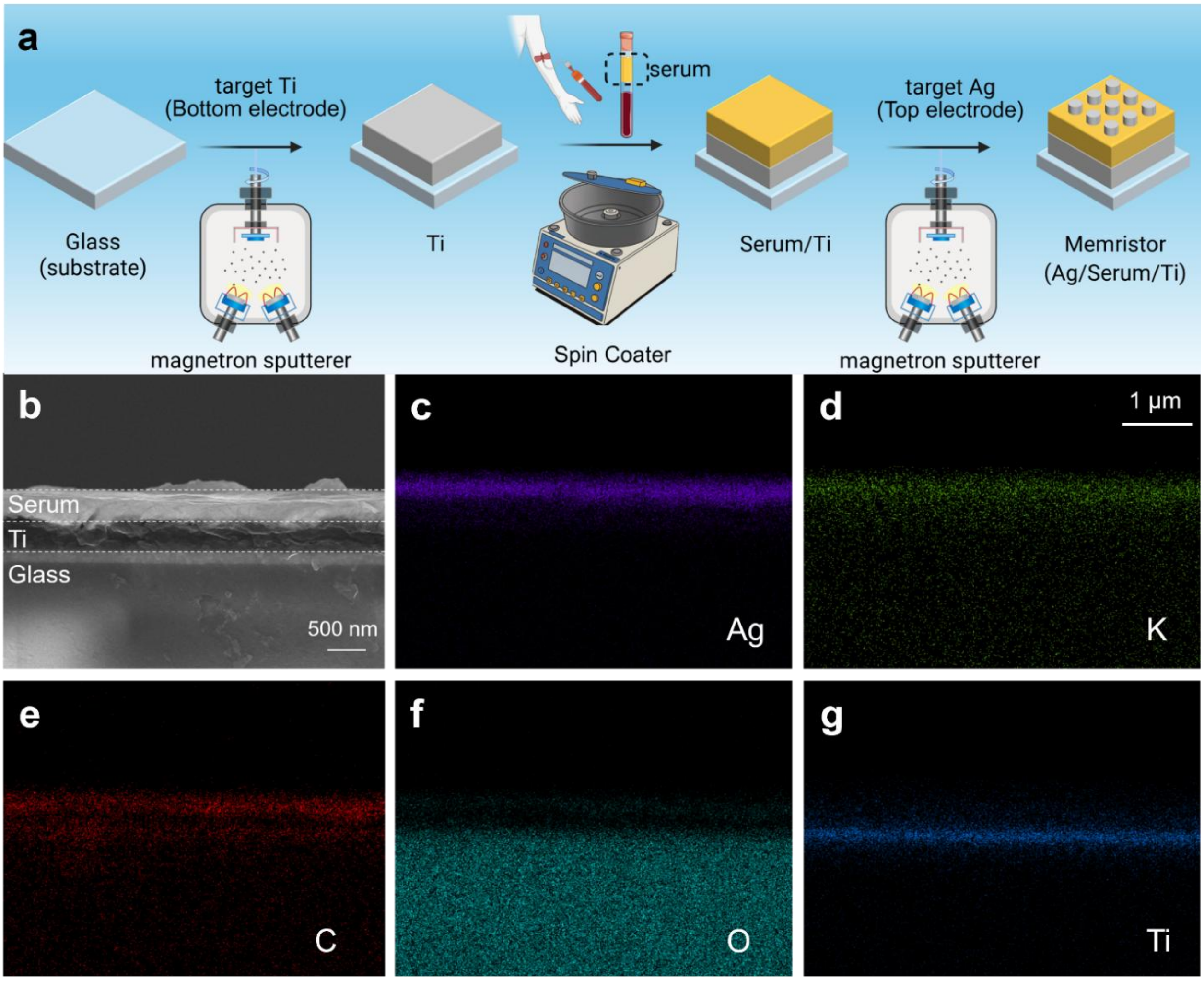


**Figure S5**. Internal characteristics the functional layer thin films of memristor with Ag/human serum/Ti structure. a) Fabrication process of memristor with Ag/human serum/Ti structure. Created with BioRender.com released under a Creative Commons Attribution-NonCommercial-NoDerivs 4.0 international license. b) SEM image of the cross-section of the memristor. c-g) Distribution of Ag, K, C, O, and Ti elements in the cross section of a memristor as observed by scanning electron microscopy (SEM).

**Supplement Text Note 5:**

Elemental distribution data for the cross-section are presented in **Figure S5**b, in which it clearly reveals that the functional layer is serum. Cross-sectional SEM-EDX analysis further visualized the elemental composition across the device layers, with Ag, K, C, O, and Ti clearly visible in color (Figure S5 c-g).

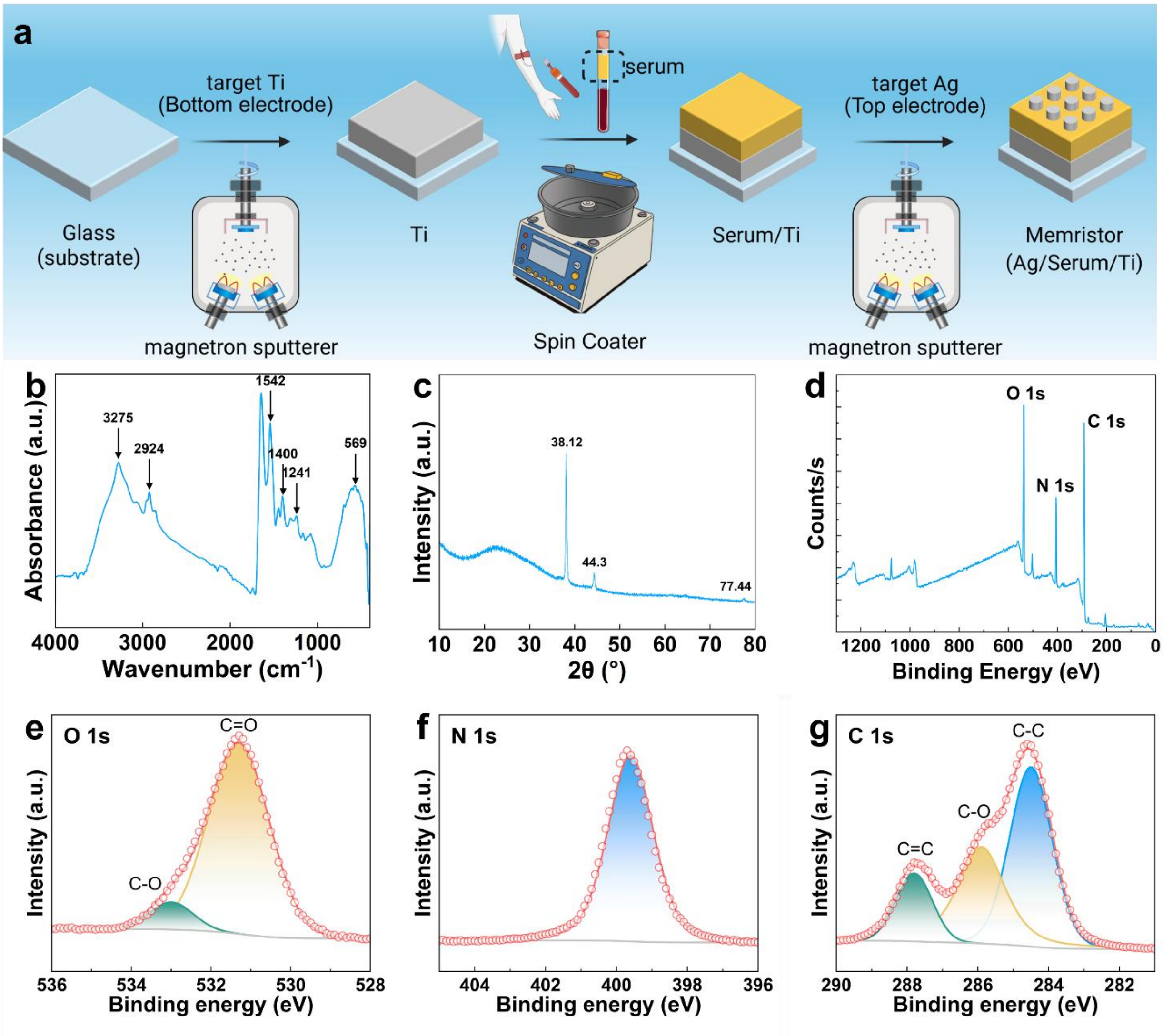


**Figure S6**. Internal characteristics of the functional thin-film layer in Ag/human serum /Ti structured memristors. a) Fabrication process of memristor with Ag/human serum/Ti structure. Created with BioRender.com released under a Creative Commons Attribution-NonCommercial-NoDerivs 4.0 international license. b) FT-IR analysis of serum. c) XRD characterization of serum. d)XPS spectrum. e) XPS test of O 1s. f) XPS test of N 1s. g) XPS test of C 1s.

**Supplement Text Note 6:**

Spectroscopic and diffraction analyses were performed to characterize the serum material properties. Fourier-transform infrared spectroscopy (FTIR, **Figure S6**b) and X-ray diffraction (XRD, Figure S6c) provided further insights into the chemical functional groups and crystalline structure of the serum, respectively. The XPS spectrum shows that the most abundant elements in the middle layer are C, N, and O (Figure S6d). High-resolution X-ray photoelectron spectroscopy (XPS) spectra of C1s, N1s, and O1s exhibited distinct binding energy peaks corresponding to these elements (Figure S6e-g), confirming their chemical states and bonding configurations within the device structure.
